# Pupil responses as indicators of value-based decision-making

**DOI:** 10.1101/302166

**Authors:** Joanne C. Van Slooten, Sara Jahfari, Tomas Knapen, Jan Theeuwes

**Affiliations:** Department of Experimental and Applied Psychology, Vrije Universiteit, Amsterdam, The Netherlands; Spinoza Centre for Neuroimaging, Royal Academy of Sciences, Amsterdam, The Netherlands; Department of Psychology, University of Amsterdam, Amsterdam, The Netherlands

## Abstract

Pupil responses have been used to track cognitive processes during decision-making. Studies have shown that in these cases the pupil reflects the joint activation of many cortical and subcortical brain regions, also those traditionally implicated in value-based learning. However, how the pupil tracks value-based decisions and reinforcement learning is unknown. We combined a reinforcement learning task with a computational model to study pupil responses during value-based decisions, and decision evaluations. We found that the pupil closely tracks reinforcement learning both across trials and participants. Prior to choice, the pupil dilated as a function of trial-by-trial fluctuations in value beliefs. After feedback, early dilation scaled with value uncertainty, whereas later constriction scaled with reward prediction errors. Our computational approach systematically implicates the pupil in value-based decisions, and the subsequent processing of violated value beliefs, ttese dissociable influences provide an exciting possibility to non-invasively study ongoing reinforcement learning in the pupil.

## Introduction

There is fast-growing interest to understand how the pupil, as a non-invasive proxy of neuromodulation^1^, relates to cognition and in particular decision-making. Traditionally, pupil dilation during and after decisions has been related to uncertainty and surprise^2-6^, likely via noradrenergic modulations by the locus coeruleus (LC)^7,8^. Recent work, however, shows that the pupil also tracks activity of other neuromodulatory nuclei. For example, in a recent perceptual decision-making task it was found that pupil dilations were modulated by activity in dopaminergic midbrain nuclei^9^, ttese nuclei are known to release dopamine in response to rewards and reward-predicting cues to optimize future decisions^10-12^, tte pupil also dilates in response to the presentation of cues predicting reward^13-15^ and tracks changes in reward expectations^16^, ttese pupil responses are blunted in Parkinson’s patients, yet are fully restored by dopamine agonists^17,18^.

These findings raise the intriguing possibility that the pupil is sensitive to a multitude of neuromodulatory processes, including dopamine, implying that it could be used to non-invasively study the underlying processes that shape value-based decisions and learning. Here, we investigated the interplay between the pupil, reinforcement learning and value-based decision-making by using a computational reinforcement learning (RL) model as a basis for linear systems analysis of pupil size fluctuations.

We measured pupil size while thirty-four participants performed a probabilistic reinforcement learning task, consisting of a learning and transfer phase (Fig. la,b & Methods), tte reliability of choice outcomes varied across three learning pairs^19^ with different reward probabilities, ttese different reward probabilities create varying degrees of choice difficulty, uncertainty and value expectations across choices. We fit a hierarchical Bayesian version of the Q-learning RL algorithm^20^ to participants’ choices in the learning phase to describe value-based choices and outcome evaluations (Fig. lc & Methods)^21-24^, tte Q-learning algorithm describes value-based decision-making using two functions: a choice function and an outcome function, tte choice function calculates the probability of choosing one option (Q-chosen) over the other (Q-unchosen), based on one’s sensitivity to value differences, or explore-exploit tendency (*β*; Fig. Id, left panel), tte outcome function then computes the magnitude by which the reward prediction error (RPE) changes value beliefs about the chosen option, scaled by the learning rate (*α*; Fig. Id, right panel)^25^. As value beliefs are differentially updated after positive and negative outcomes^26-28^ via different striatal learning mechanisms^29-31^, we defined separate learning rate parameters for positive (*β_Gain_*) and negative (*α_Loss_*) choice outcomes^21,26,27,32^.

**Figure 1:**
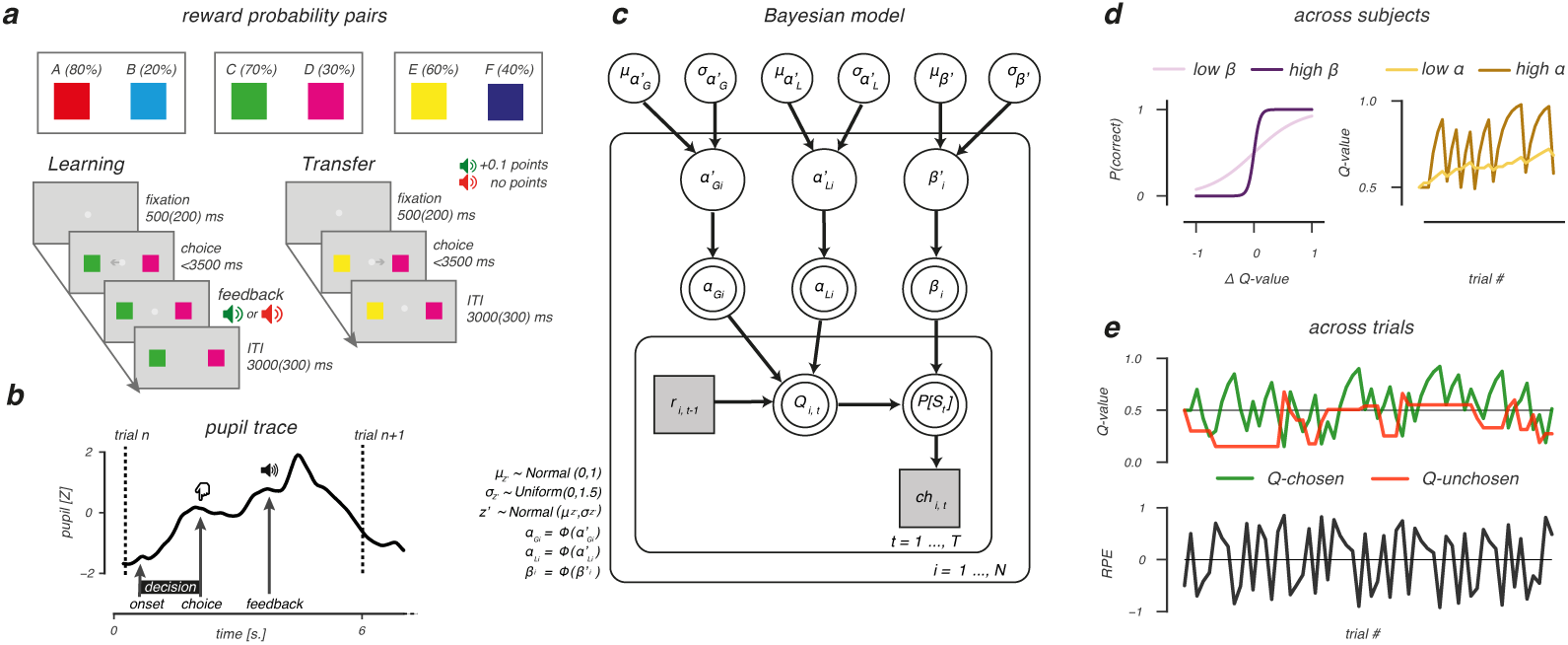
Probabilistic selection task and reinforcement learning model. **a**, During learning, 3 option pairs were presented in random order. Participants had to select the more rewarding option of each pair (option A, C and E) by learning from probabilistic feedback that indicated +0.1 reward points after a ‘correct’ choice, or no points. Choosing option A resulted in a reward in 80% of the times, whereas choosing option B resulted in a reward only in 20% of the times. Reward probability ratios were 70/30 for the CD pair and 60/40 for the EF pair, thereby increasing uncertainty about the correct option to choose, tte transfer phase tested how much was learned from the probabilistic feedback. All options were randomly paired with one another, and participants selected the most rewarding option based on earlier learning. In this phase, feedback was omitted, **b**, Example pupil trace for a trial in the learning phase, **c**, Bayesian hierarchical model, consisting of an outer participant (*i* = l…,*N*) and inner trial (*t* = 1…,*T*) plane. Variables of interest are depicted by circular and squared nodes, indicating continuous and discrete variables, respectively. Shaded variables are obtained from the behavioural data and used to fit the model. Double bordered variables are deterministic, as they were derived from the model fit. Arrows indicate dependencies between variables. Φ() represents the probit transform, **d**, Model parameters governing value-based decision-making. Left panel: the *β*-parameter describes sensitivity to option value differences (ΔQ-value). Higher *β*-values indicate greater sensitivity to ΔQ-value and more exploitatory decisions for options with highest expected rewards. Right panel: the *α*-parameter governs value belief updating. Higher learning rates (*α*) indicate rapid, but more volatile value belief updating compared to lower learning rates, **e**, Across-trial fluctuations in value beliefs (Q-values) for the chosen and unchosen option and RPEs with the EF pair as example.

Our computational approach allows us to investigate the potential utility of the pupil as a proxy for value-based decision-making and value belief updating, across two levels. First, we describe participants’ choice behaviour using parameters that embody core computational RL principles, ttese parameters provide a strong handle to investigate how inter-individual differences in value-based learning and decision-making relate to pupil responses. Second, by simulatingthe learning process we can investigate how pupil size depends on trial-to-trial fluctuations in underlying computational variables such as value beliefs, uncertainty and reward prediction errors, ttat is, our experimental paradigm allowed us to map pupil responses onto separable computational components both across participants and trials.

## Results

### Behavioural and model performance

Participants learned the stimulus-reward contingencies well, as they correctly learned to select the higher reward probability option in all three pairs (*P*(correct) above chance, all *P*’s <.001; Fig. 2a). Performance was best in AB and decreased progressively from CD to EF, where smaller differences in the reward probability ratios increased the number of incorrect responses (*F*_(2,66)_ = 14.45, *P*<.001,
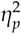
=.19) and response times (*F*_(2,66)_ = 5.5, *P*=.006,
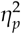
=.04). In the transfer phase, choices were guided by the previously learned reward probabilities. Here, participants made more errors (E(2,66)= 49.3, P<.001, 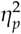=53) and were slower (*F*_(2,66)_=34.6, P<.001,
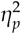
=.12)
when confronted with option pairs with small value differences (Fig. 2b), consistent with earlier studies.^26,33,34^

**Figure 2:**
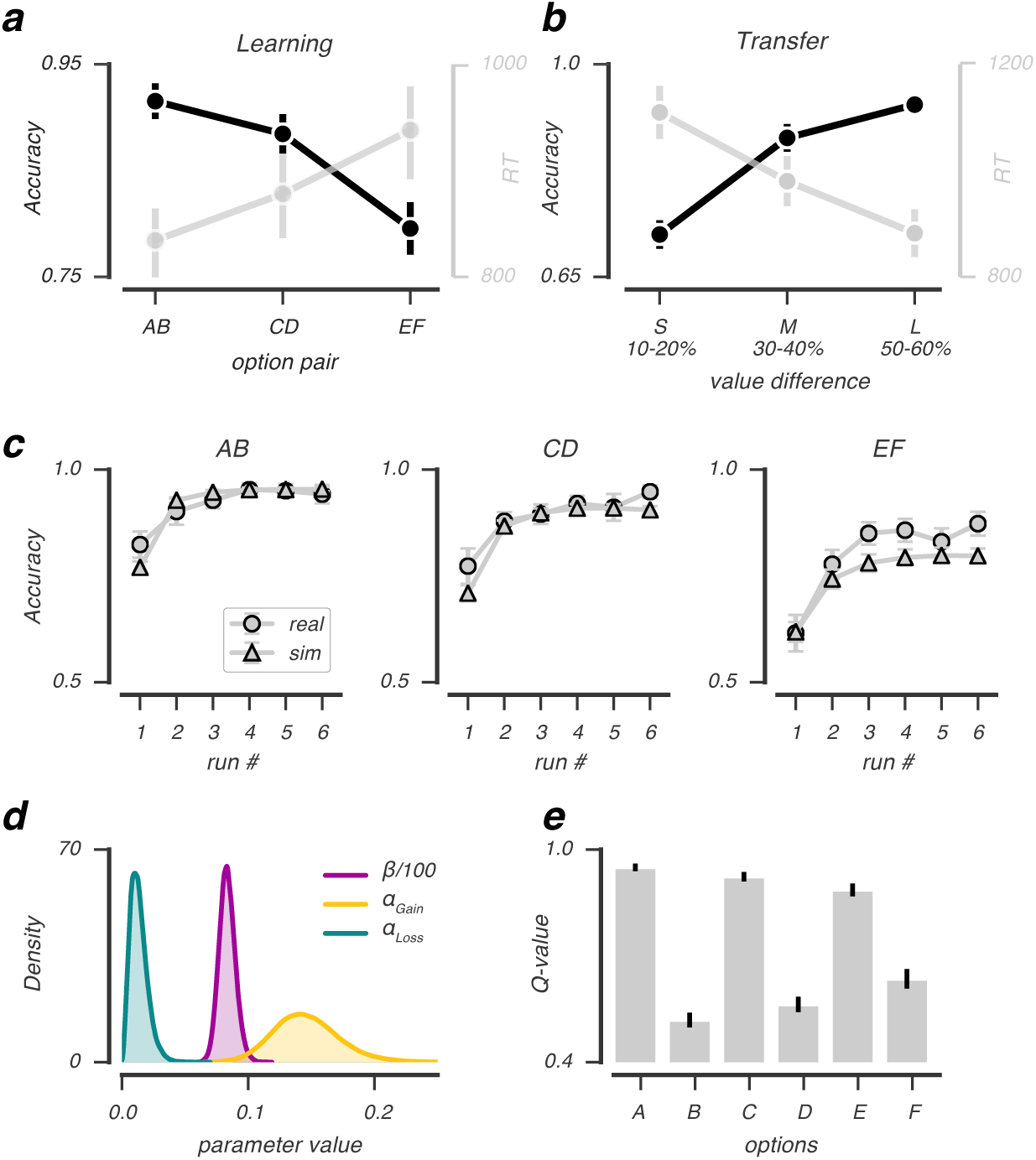
Behavioral and model performance. Average accuracy and RT across subjects (N=34) as a function of option pairs in the learning phase (**a**) and option value differences (derived from the experimental reward probabilities) in the transfer phase (**b**) that indicated small (S), medium (M) or large (L) value differences between presented options, **c**, Real and simulated choice accuracy as a function of run number in the learning phase, split by option pair. For all option pairs, simulated and real accuracy was very similar, with real EF accuracy being slightly underestimated by the model, **d**, Group-level posterior distributions of the obtained parameter estimates for *β*, *α_Gain_* and *α_Loss_*. **e**, Model estimates of value beliefs for each option at the end of the learning phase, *β*/100 for visualization; error bars represent mean ± s.e.m.

The Q-learning model simulated participants’ choice behavior well (Fig. 2c) when using the fitted learning rates (*α_Gain_*, *α_Loss_*) and explore-exploit (*β*) parameter (Fig. 2d). In accordance with behavior, the estimated value beliefs were highest for A and lowest for B (Fig. 2d) with differences in value beliefs being largest for AB, followed by the CD and EF pair (*F*_(2,66)_= 20.63, *P*<.001,
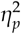=.39).

### Pupil responses predict individual differences in value-based decision-making

We next investigated whether the pupil was sensitive to the underlying processes supporting value-based decisions. To do so, we first characterized the average pupil response pattern across subjects epoched around two separate moments in the trial: leading up to and immediately after the moment of choice and after receiving choice feedback. Around the moment of a choice, average pupil dilation was observed already ∼ls. prior to the moment of the behavioural report (Fig. 3a), reflecting the unfolding decision process^5,35^. After receiving choice feedback, a biphasic pupil response was observed that was characterized by early dilation (∼ls. post-event) and late constriction (∼2s. post-event; Fig. 3b).

**Figure 3:**
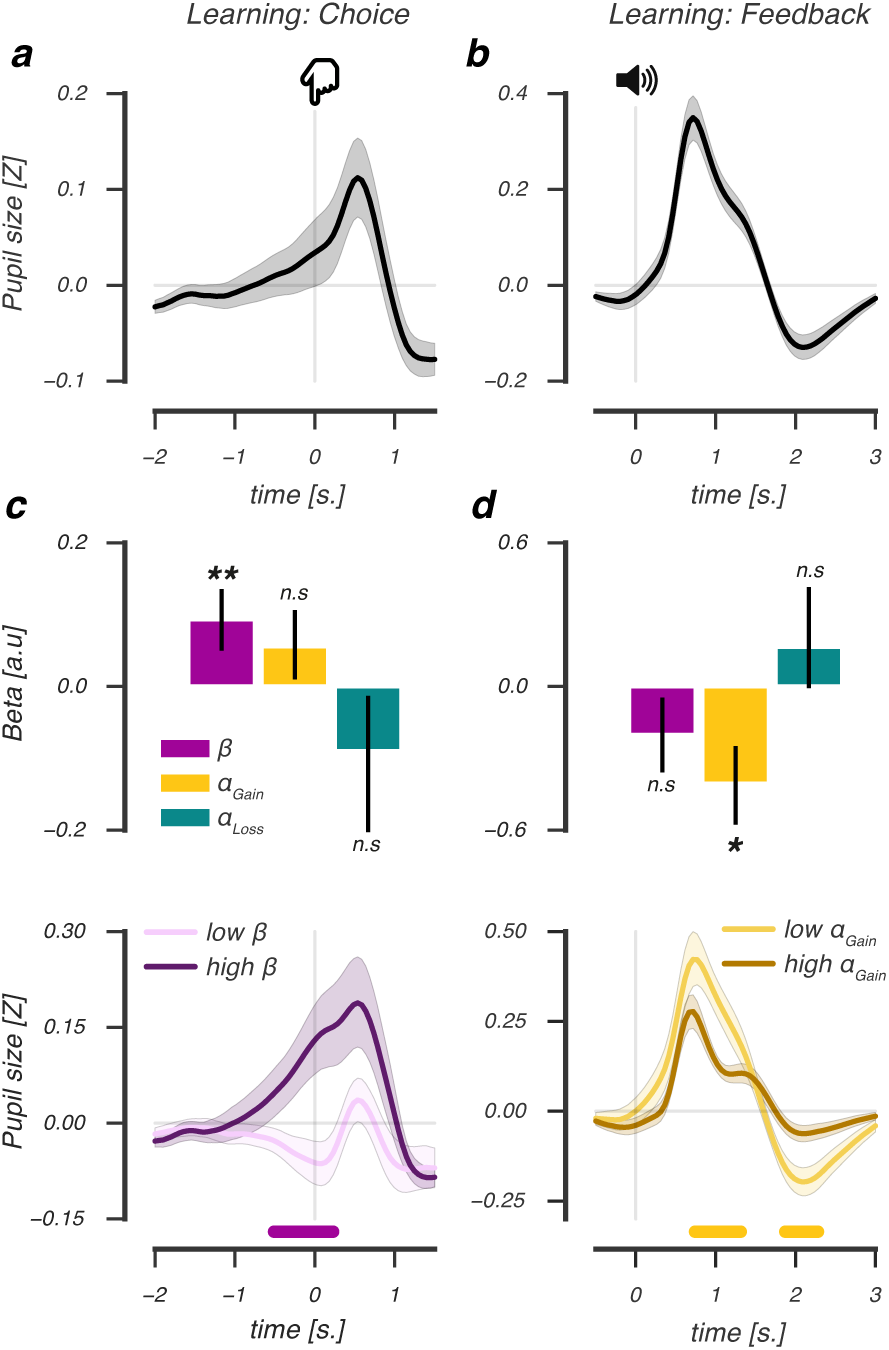
Across-subject relations between model parameters and pupil responses during choice and after feedback. Average deconvolved choice-related (**a**) and feedback-related (**b**) pupil response. Regression coefficients of an across-subject GLM of the relation between derived model parameters and pupil dilation at the moment of choice (**c**, upper panel), and a scalar amplitude measure of the feedback-related pupil response (**d**, upper panel). Median split across subjects based on modulations of *β* at the time of choice (**c**, lower panel), and *α_Gain_* after feedback (**d**, lower panel). Lines and (shaded) error bars of represent mean ± s.e.m of across-subject modulations (N=34). Horizontal significance designators indicate time points where regression coefficients significantly differentiate from zero (P<.05), based on cluster-based permutation tests (n=1000), ^⋆⋆^*P*<.01, ^⋆^*P*<.05.

Across individuals, the observed choice- and feedback-evoked pupil responses corresponded differentially to the underlying processes driving value-based decision-making. As shown in Fig. 3c (upper panel), pupil dilation at the moment of a choice was uniquely predicted by an individual’s sensitivity to value differences, or explore-exploit tendency (*β*; permutation test, *p*=.006; Supplementary Fig. la), indicating that a greater tendency to exploit high value options (high *β*) related to a stronger dilatory response (Fig. 3c, lower panel). Feedback-related dilation and constriction correlated inversely with an individual’s positive, but not negative, learning rate (Supplementary Fig. lb), suggesting that this parameter selectively scaled the amplitude of the feedback-evoked pupil response. Indeed, as shown in Fig. 3d (upper panel), the feedback-evoked response amplitude was uniquely predicted by an individual’s positive learning rate (*α_Gain_*; permutation test, *P*=.017), indicating that slower updating of value beliefs after positive feedback predicted a stronger feedback-evoked response (low *α_Gain_*; Fig. 3c, lower panel & Supplementary Fig Id).

In sum, choice- and feedback-evoked pupil responses differentially predicted the underlying processes supporting value-based decisions in the learning phase, tte tendency to exploit high value options (*β*) predicted stronger pupil dilation leading up to a value-driven choice, whereas less updating of value beliefs after positive feedback (*α_Gain_*) predicted an amplified feedback-related response, ttese relations are consistent with theories that describe and formalize Q-learning, in which the explore-exploit parameter determines the outcome of a value-driven choice and learning rates affect how much value beliefs are updated after receiving choice feedback.

### Pupil dilation reflects the value of the upcoming choice, during learning but not in transfer

We observed that across-subject variability in pupil responses was explained by model parameters that describe the underlying processes driving value-based decision-making. But do pupil responses also reflect the ongoing reinforcement learning process during value learning? In a next step, we in-vestigated the extent to which trial-to-trial fluctuations in variables describing ongoing value-based decision-making were tracked by pupil responses.

In the learning phase, prior to reaching a value-driven choice, pupil dilation correlated positively with the value difference between options (cluster *P*<.001, 2.0s. pre-event until −0.07s. pre-event, Fig. 4a, upper panel), indicating that larger value differences elicited larger pupil dilation before the choice. Specifically, the pupil dilated as a function of trial-by-trial value beliefs of the chosen, but not the unchosen option (paired *t*-test, *t*(33)=6.98, *P*<.001; Fig 4b, upper panel), revealing that pupil dilation uniquely reflected the value belief determining the upcoming choice.

**Figure 4:**
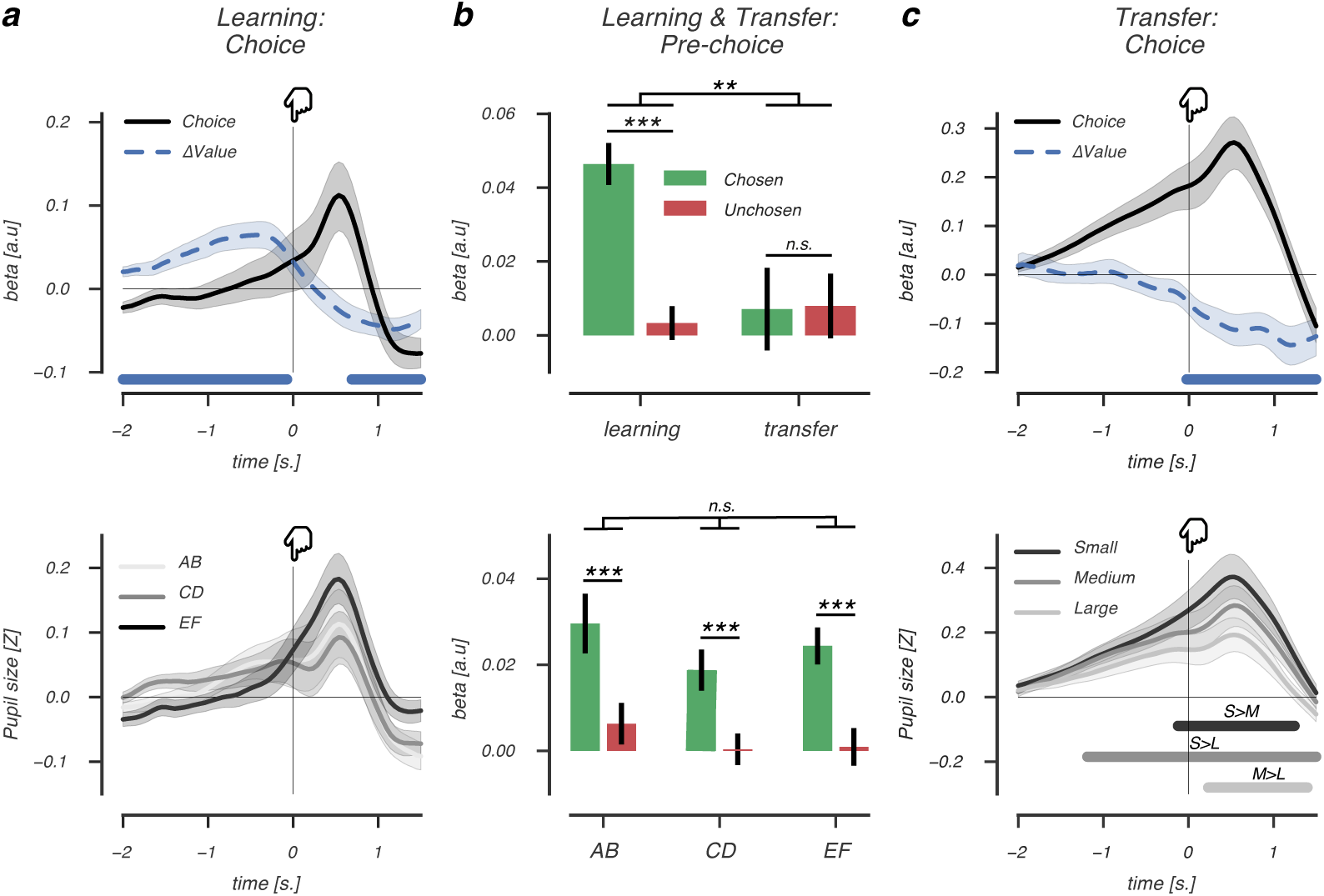
Pre-choice pupil dilation reflects the value of the upcoming choice. **a** (upper panel), Beta coefficients accounting for choice-related pupil dilation in the learning phase. Larger value differences between options (blue dashed line) elicited larger choice-related pupil dilations (black solid line) prior to choice (at *t*=0). After choice, this relationship reversed, as smaller value differences elicited larger post-choice pupil dilations, **a** (lower panel), Average choice-related pupil dilation for AB, CD and EF pairs, **b** (upper panel), Beta coefficients of chosen and unchosen value regressors accounting for pupil size fluctuations in the pre-choice decision interval of the learning (left) and transfer phase (right), **b** (lower panel), Beta coefficients of chosen and unchosen value regressors split by learning phase pairs, showing that pre-choice pupil size is modulated by values of the to-be chosen stimulus, irrespective of uncertainty, **c** (upper panel), Learned value differences did not modulate choice-related pupil dilation prior to choice (at *t*=0). After choice, smaller learned value differences elicited stronger pupil dilation, **c**, (lower panel): Smaller value differences between choice options elicited larger post-pupil dilation, indicating choice conflict drove pupil size. Lines and (shaded) error bars of represent mean ± s.e.m of within-subject modulations. Horizontal significance designators indicate time points where regression coefficients significantly differentiate from zero (*P*<.05), based on cluster-based permutation tests (n=1000), ^⋆⋆⋆^ *P*<.001, ^⋆⋆^ *P*<.01, repeated measures ANOVA.

To rule out the possibility that condition differences (i.e. AB, CD, EF) instead of trial-by-trial fluctuations in chosen value beliefs explained pupil dilation prior to a choice, we estimated their independent effects on pupil size in a single regression analysis. We observed no differences between conditions in average pupil dilation prior to a choice (Fig. 4a, lower panel), ttis also excluded the hypothesis that pre-choice pupil dilation was driven by uncertainty, as we did not observe significantly more dilation in the most difficult, hence most uncertain, EF pair. In all pairs, pre-choice pupil size correlated positively with chosen value (*F*_(2,66)_ = 19.76, *P*<.00l,
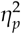
=.15; Fig. 4b, lower panel) irrespective of the condition type (*F*_(2,66)_ = 1.8, *P*=.17). ttus, prior to reaching a value-driven choice, the pupil tracked subtle differences in value beliefs about the upcoming choice, while dilation was not driven by uncertainty differences between the conditions.

Next, we asked whether value beliefs also modulated pre-choice pupil dilation in the transfer phase, where feedback was omitted. In contrast to the learning phase, pupil dilation prior to a value-driven choice was not predicted by previously learned value differences between options (Fig. 4c, upper panel), nor by separate chosen or unchosen value beliefs (paired *t*-test, *t*<l, Fig. 4b, upper panel). Indeed, a repeated measures ANOVA with the factors phase (learning, transfer) and value (chosen, unchosen) indicated that only during learning, but not during transfer, pre-choice pupil dilation was modulated by value beliefs about the upcoming choice (*F*_>33_)= 6.9, *P*=.013,
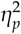
=.06).

However, immediately after a value-based choice, learned value beliefs negatively predicted pupil dilation both in the learning (cluster *P*=.007, 0.68s. pre-event until 1.5s. post-event; Fig. 4a, upper panel) and transfer phase (cluster *P*=.003, −0.02s. until 1.48s. post-event; Fig. 4c, upper panel). Now smaller, instead of larger, value differences elicited larger post-choice pupil dilation, suggesting that the difficulty of a recent choice, or the choice conflict it generated, drove pupil size upward. Indeed, we observed a similar post-choice pupil response pattern when regressing choice conflict on the basis of the experimental reward probabilities on pupil size (Fig. 4c, lower panel), indicating that post-choice pupil dilation was modulated by choice conflict, consistent with an earlier report^34^.

These model-based trial-to-trial analyses show that when engaged in active reinforcement learning, pupil dilations differentially reflect value beliefs and choice conflict at different points in time. Prior to value-based choices, pupil size uniquely reflected value beliefs about the upcoming choice, where stronger dilations predicted higher value beliefs, ttis pattern of pre-choice value dilations was absent in the subsequent transfer phase where rewards could not be obtained, indicating that seemingly similar pupil dilations prior to value-based choices indexed different cognitive processes during learning and transfer.

### Feedback-evoked pupil responses reflect value uncertainty and reward prediction errors

Only during active reinforcement learning, we observed that choice-related pupil dilation reflected value beliefs about the upcoming choice. If the pupil reliably tracked ongoing reinforcement learning, it should also provide information about the evaluation of a choice outcome. In the last step, we therefore investigated how feedback-evoked pupil responses covaried with the degree to which outcomes violated value beliefs about a recent choice.

We observed larger feedback-evoked pupil dilation (Fig. 5a) after choices between options with small value differences. Specifically, early post-feedback dilation correlated negatively with differences in value beliefs of recently presented options (cluster *P*<.001, −1.5s. pre-event until 1.78s. post-event; Fig. 5b). We furthermore verified that these feedback-evoked dilations were not driven by feedback valence (Supplementary Fig. 2a,b). In contrast to dilation in the choice interval, dilation in the feedback interval was explained by fluctuations in trial-by-trial value beliefs of both the chosen and the unchosen options, in opposite directions (Fig. 5c). ttus, lower beliefs about the chosen and higher beliefs about the alternative option both increased dilation, indicating that uncertainty about the value of a past choice modulated feedback-evoked dilation. In support of this, trial-by-trial chosen and unchosen value beliefs explained feedback-evoked dilation already prior to receiving feedback, showing that it was uncertainty about the outcome of a value-based decision that drove pupil size. Lastly, outcomes that violated value beliefs did not elicit larger feedback-evoked dilations (Supplemental Fig. 2), excluding the hypothesis that these modulations of the feedback response reflected surprise.

**Figure 5:**
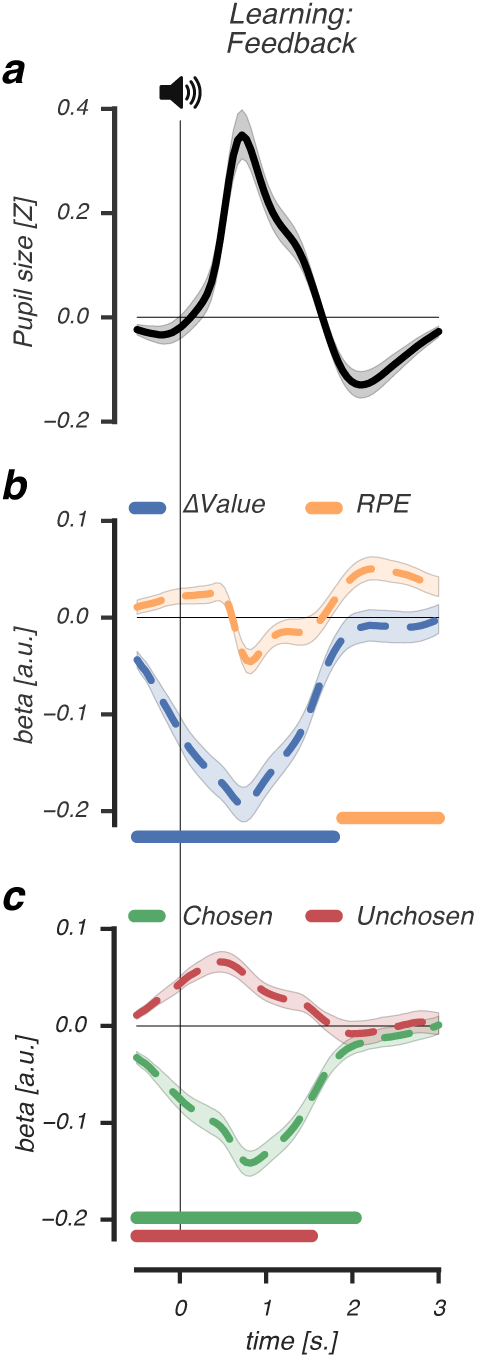
Feedback-evoked pupil responses reflect value uncertainty and reward prediction errors. **a-c**, Beta coefficients accounting for feedback-evoked pupil responses in the learning phase, tte feedback-evoked pupil response (**a**) was characterized by early dilation (∼ls. post-event) and late constriction (∼2. post-event), **b**, Early in time (∼ls. post-event), feedback-evoked pupil dilation correlated negatively with the difference in value beliefs about recently presented options (ΔValue, blue dashed line). Late in time (∼2s. post-event), feedback-evoked pupil constrictions correlated positively with signed RPEs (orange dashed line), **c**, Both lower value beliefs of the recently chosen and higher value beliefs of recently unchosen option increased feedback-evoked pupil dilations, already prior to the moment of feedback (at *t*=0). Lines and shaded error bars represent mean ± s.e.m of within-subject modulations. Horizontal significance designators indicate time points where regression coefficients significantly differentiate from zero (*P*<.05). Statistics based on cluster-based permutation tests, n=1000.

Importantly, whereas value beliefs about a recent choice affected early dilation, the degree to which outcomes violated those beliefs modulated late feedback-evoked pupil constriction. As shown in Fig. 5b, signed RPEs correlated positively with late feedback-evoked pupil constriction ∼2s. after receiving feedback (cluster *P*<.00l, 1.8s. until 3.0s. post-event), ttis correlation indicated that worse-than-expected outcomes (-RPEs) elicited stronger pupil constriction compared to better-than-expected out-comes (+RPEs).

To summarize, we observed a biphasic feedback-evoked pupil response that tracked the evaluation of a recent value-based choice. Early pupil dilation was modulated by uncertainty about the value of options, as choices between similarly valued options increased dilation the most. Late pupil constriction was modulated by the violation of current value beliefs, as worse-than-expected outcomes elicited stronger pupil constriction compared to better-than-expected ones.

## Discussion

The present results provide the novel insight that the pupil is a reliable reporter of the underlying process of learning and decision-making based on value. When engaged in active reinforcement learning, but not when choice value was already internalized, the pupil showed two distinct response patterns. Prior to reaching a value-driven choice, pupil dilations scaled with trial-by-trial value beliefs about the upcoming choice and were diagnostic for an individual’s sensitivity to choose the option with the highest expected outcome. Feedback about the choice subsequently evoked a biphasic evaluation response. Early pupil dilation scaled with uncertainty about the value of recent choice options, whereas subsequent pupil constriction scaled with the violation of current choice value beliefs, or signed reward prediction errors. Moreover, the amplitude of this (post-feedback) biphasic response was predicted by variability in learning rates across participants, which determine the updating of value beliefs given the reward prediction error.

Previous studies have shown that cues predicting reward increase pupil dilation^13-15,17,18^. Our observation that value beliefs increase pupil dilation prior to choice are in line with these findings. Critically, we additionally show that pupil dilation increases as a function of value beliefs about the chosen option and signal the value of the upcoming choice. Moreover, we found that pupil dilations were not modulated by the value of the alternative option, indicating that the pupil does not reflect the values associated with all potential options, but specifically, the value that was driving the choice. Interestingly, striatal dopamine concentrations and phasic responses of dopamine neurons are observed to reflect chosen value^36-38^, ttis signalling is thought to support value learning of a choice, as the value of the chosen option gets updated according to the reward prediction error after receiving choice feedback.

Importantly, only during learning, but not during transfer, was pupil dilation prior to choice modulated by choice value. Why was this the case, when participants had to make value-based decisions in both phases? One important difference between the phases was the ability to learn from the outcomes of actions. In the learning phase, options could be compared to each other and the outcome of a choice was immediately presented, ttis was very different in the transfer phase, when participants were confronted with new choice situations and had no ability to learn from their choices because they never received feedback. Dopamine, particularly in the striatum, plays and important role during reinforcement learning^39^. Striatal dopamine strengthens actions that lead to rewarding outcomes and weakens those that lead to aversive ones^19,26,30^, thereby flexibly adapting behaviour to maximise rewards. In the transfer phase, value beliefs are consolidated and dopamine no longer plays an important role in learning or modulating choice behaviour. Information used to make a value-based choice is now retrieved from memory, guided by structures that encode learned value representations, such as the ventromedial prefrontal cortex^40,41^. Our finding that the pupil was only sensitive to choice value during active reinforcement learning could mean that the pupil is particularly sensitive to the process of learning the value of choices, or contingency learning.^42^

An individual’s sensitivity to value differences between presented options, as quantified by our computational model, predicted the amount of pupil dilation exactly at the time of a value-based choice. Individuals that were more sensitive to small value differences showed stronger dilations, ttese individuals exploited high value options, leading to better task performance^22^. Increased pupil dilation has been associated with better task performance^43,44^, or with the tendency to exploit in dynamic environments^45,46^, tte observed relationship between pupil dilation and individual sensitivity to value differences could reflect either of these processes, as choosing a high value option can result from accurate option value representations or from the general tendency to favour exploitation over exploration^47^. Measuring pupil responses during value-based decision-making in a reversal learning paradigm might provide a way to disentangle these two alternative explanations, as optimal task performance would then depend on changing decision strategies over time.

Uncertainty did not drive pupil size prior to a value-based choice, as our results indicated that the most difficult, hence uncertain, pair (EF) did not increase dilation. On the contrary, higher value beliefs, indicating more certainty about choice value, elicited greater dilation, ttis finding contrasts with earlier work relating pupil dilations to situations of increased uncertainty^2,4,5^, but aligns with those associating it with certainty^3,48^. One possible explanation for these differential findings is that in our study, higher value beliefs about the upcoming choice led to increased reward expectations^17^, or decreased risk assessment^3^, driving pupil dilation prior to the choice.

However, immediately after a value-based choice, but prior to feedback, we observed that uncertainty modulated pupil size, as choices between closely valued options triggered strong post-choice dilation. While our study is the first to relate the pupil directly to value beliefs, these findings are consistent with observations of pupil dilation indexing choice conflict after difficult decisions^34^ or decision uncertainty during perceptual decisions^6^. Our work extends these studies by showing that different cognitive processes affect pre- and post-choice dilation, linking pre-choice dilation to chosen value beliefs and post-choice dilation to value uncertainty, ttis provides a tentative link to dopamine neurons encoding reward uncertainty^49^, where sustained dopamine activity prior to the moment of the outcome was strongest after cues predicting reward with 50% probability, ttese highly unpredictable cues are thought to drive learning maximally, as high subjective uncertainty indicates the lack of an accurate reward predictor and the need to improve predictions^49^. Pupil dilation after a choice between options with small value differences may relate to this process, reflecting the allocation of attentional resources to support learning from the upcoming choice outcome.

After feedback, a biphasic feedback-evoked pupil response tracked two different evaluation processes related to the outcome of a choice. First, early dilations were explained by uncertainty about the value of the choice outcome, but not by surprise, as outcomes that violated value beliefs did not trigger stronger dilations, ttis is a somewhat surprising finding, given that pupil dilations can be stronger after unexpected outcomes^3,4,50-52^. What our results suggest is that during reinforcement learning, the uncertainty associated with the outcome of a choice increases pupil dilation, irrespective of its positive or negative value. What this may indicate is that early pupil dilation reflected increased attention to uncertain, but potentially rewarding stimuli in the environment.

Second, late pupil constriction was explained by signed reward prediction errors, thus, reflecting how much the outcome violated current value beliefs about the chosen option. Lower-than-expected choice outcomes resulted in smaller pupil responses compared to higher-than-expected ones, which shows a striking resemblance to the reward prediction error pattern of phasic dopamine neurons^10,53,54^ that briefly activate after higher-than-expected outcomes and deactivate after lower-than-expected outcomes. In calculating this error term, the obtained reward is compared to the value of the chosen option^36^, which we observed also modulated pupil dilation prior to a value-based choice, ttus, our findings suggest that pupil responses track specific decision and evaluation signals that promote value-based learning and decision-making.

Recently, a more elaborate view of the phasic dopamine reward prediction error response has been proposed, in which an initial unselective component detects any potential reward and a later component codes the well-known reward value^55^ tte dynamic evolution of the feedback-evoked pupil response is consistent with this pattern, as early dilation unselectively increased after uncertain outcomes and late constriction reflected the evaluation of the outcome’s reward value.

To conclude, our study provides evidence that the pupil is a reliable indicator of value-based decision-making, as it signalled the processing of value up to a choice and the subsequent evaluation of choice outcomes in terms of uncertainty and violations of value beliefs, ttere were several aspects to our approach that enabled us to establish these specific relations and to move beyond previous work linking the pupil to reward. First, we characterised the full temporal profile of value-based decisions in the pupil, thereby relating different decision and evaluation processes to different components of the pupil response. Second, these specific relations could only be established with the use of a formal learning model that provided us access to participants’ developing value beliefs, and the underlying processes thought to support value-driven decisions, ttis also highlights our use of subjective value estimates to relate to the pupil. In contrast, previous studies investigating reward-related effects on pupil size employed externally defined value estimates^2,3,17,18,52^, with one notable exception^16^. Lastly, our study describes the temporal evolution of reinforcement learning in the pupil, thereby providing evidence that the pupil can be used to non-invasively track the reinforcement learning process as it takes place. Future studies that combine functional brain imaging and pupillometry will have to further specify the brain areas that contribute to the value-based pupil response.

## Methods

### Participants

Forty-two healthy participants with normal to corrected to normal vision completed the experiment (10 males; mean age=24.9; age range=18-34 years), ttey were paid 16€ for 2 hours of participation and earned an additional performance bonus (mean=10.2€, SD=1.8). tte ethical committee of the Vrije Universiteit Amsterdam approved the study and written informed consent was obtained from all participants. Eight participants were excluded from analyses due to the following reasons: inadequate fixation to the center of the screen (N=4), reporting more than three unique stimulus pairs in the learning phase (N=l) and (almost) perfect choice accuracy in the learning phase, which complicated behavioral model fitting (N=3), resulting in a total of 34 participants for the analyses.

### Task & Procedure

Participants were seated in a dimly lit, silent room with their head positioned on a chin rest, 60 centimeters away from the computer screen, ttey received written information about the general purpose of the experiment, after which they completed a 30-trial practice session of the learning phase. Subsequently, participants completed for the learning phase 6 runs of 60 trials each (360 trials in total, 120 presentations of each stimulus pair), with small breaks in-between runs. After each run, the earned number of points was displayed. At the end of the learning phase, the total number of earned points was converted into a monetary bonus. Directly after the learning phase, participants entered the transfer phase, ttey completed 5 runs of 60 trials each (300 trials in total, 20 presentations per stimulus pair), with small breaks in-between runs. Overall choice accuracy was displayed at the end of the transfer phase.

### Stimuli & trial structure

Stimuli were presented on a 21-inch Iiyama Vision Master 505 MS103DT with a spatial resolution of 1024 × 768 pixels, at a refresh rate of 120Hz, with mean luminance 60 cd/m2. Experiments were programmed in OpenSesame and data analysis were performed using custom software written in Python, using Numpy (vl.11.2), Scipy (vO.18.1), FIRDeconvolution (vO.l.devl), Hedfpy (vO.O.devl), MNE (v0.14) and PyStan (v2.14) packages. Luminance effects on pupil size were minimized by keeping the background luminance of the display constant. Color stimuli were near-isoluminant to each other and the background (set via a flicker-fusion color calibration test carried out once at the start of the experiment). To account for luminance bias effects, each participant had a unique color pair (red-blue; yellow-dark blue; green-magenta) to reward probability mapping (AB, CD, EF) that was counterbalanced in order (e.g. red-blue or blue-red for AB).

In each learning phase trial, participants continuously fixated on a central white fixation dot. After 500ms (SD=200ms), two colored stimuli (1.26°×l.26° visual angle) appeared at the horizontal meridian left and right from the central fixation dot at a distance of 5.04° visual angle. Participants made a choice for one of the options using the ‘K’ (left choice) and ‘L’ (right choice) keys. A choice was highlighted by a small dark grey arrow (150ms) pointing in the direction of the chosen option. After a random interval drawn from a Gaussian distribution with a mean of 1500ms (SD=300ms), the choice was followed by auditory feedback, indicating reward (+0.1 points; 500ms ‘correct’ sound) or no reward (500ms; pure sine tone at 300Hz). Omissions or response times (RTs) longer than 3500ms were followed by a neutral tone (500ms; pure sine tone at 660Hz). Inter-trial intervals were drawn from a Gaussian distribution with a mean of 3000ms (SD=300ms) Trials of the transfer phase followed the same trial structure as trials in the learning phase, but had a shorter duration as choices were not followed by feedback.

### Behavioural analysis

Choices and RTs were recorded for all trials in the learning and transfer phase. RT on every trial was computed as the time from onset of the stimulus pair until the choice (keypress). Trials with RTs below 150ms or above the RT deadline of 3500ms were removed from all analyses. As a choice between two options in the learning phase was never necessarily “correct”, we defined the selection of the optimal option (more reinforcing option of the presented pair) as a correct choice. For the transfer phase, value conflict on a particular trial was defined on the basis of the experimental reinforcement value difference between the presented stimuli, where smaller value differences were associated with higher conflict.

### Computational model

Choices during the learning phase were fit with a reinforcement learning (“Q-learning”) model^20,56^. For each option, the model estimates its expected value, or “Q-value”, on the basis of individual sequences of choices and outcomes. All Q-values were set to 0.5 before learning. After each choice, the chosen option’s Q-value is updated by learning from feedback that resulted in an unexpected outcome, which is captured by the RPE, *r_i_*(*t*)-*Q*_i_(*t*). Thus, the Q-value for option *i* on the next trial *t* is updated depending on the outcome, *r*, using the following formula:

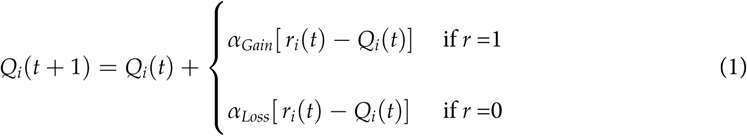

where parameters 0 ≤ *α_Gain_ α_Loss_* ≤ 1 represent positive and negative learning rates, respectively, that determine the magnitude by which value beliefs are updated depending on the RPE. We modeled separate learning rates, as different striatal subpopulations are involved in positive and negative feedback learning^29–31^ and individuals tend to learn more from positive feedback^21,22,28^. Given the Q-values, the probability of selecting one option over the other (e.g. selecting option A over B) was described by a softmax choice rule:

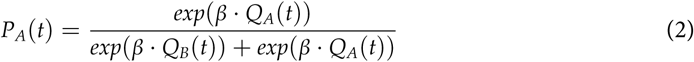

Here, 0 ≤ *β* ≤ 100, or the explore-exploit parameter, described the sensitivity to option value differences, where larger *β* values indicates greater sensitivity, and more exploitative choices, for options with relative higher reward values.

### Bayesian hierarchical fitting procedure

The Q-learning model was fit using a Bayesian hierarchical fitting procedure, where individual parameter estimates were drawn from group-level parameter distributions that constrained the range of possible individual parameter estimates, ttis procedure allowed for the simultaneous estimation of group-level and individual-level parameters^23,57^, thereby capitalizing on the statistical strength offered by the degree to which participants are similar with respect to the model parameters as well as taking into account individual differences^58^.

As shown in Fig lc, our model was implemented following Jahfari et al. (2016,2018). Variables *r_i_*(*t*-1) (outcome for participant *i* on trial *t*-1) and *ch_i_*(*t*) (choice of participant *i* on trial *t*) were obtained from the behavioural data. Per-participant parameter estimates *α_Gi_* (*α_Gain_* participant *i*), *α_Li_* (*α_Loss_* participant) and *β_i_* (*β* participant *i*) were modeled using a probit transformation
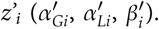
The probit transformation is the inverse cumulative distribution function of the normal distribution that can be used to specify a binary response model.
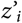
were drawn from group-level normal distributions with mean *μ_z′_*, and standard deviation *δ_z′_*. A normal prior was assigned to group-level means *μ_z′_*∼ 𝒩( 1,0) and a uniform prior to the group-level standard deviations *δ*_z’_∼𝒰( 1,1.5)^23^. tte Bayesian hierarchical model was implemented in STAN^59^ and fit to all trials of the learning phase that fell within the correct response time window 150ms ≤ RT ≤ 3500ms,(mean=99.5% of trials, SD=0.8%). Multiple chains were generated to ensure convergence, which was evaluated by the Rhat statistic^60^, tte Rhat statistic confirmed convergence of the fitting procedure (i.e., all Rhats were equal to 1.0). We also tested whether the derived per-participant parameters could simulate choices that were qualitatively similar to the observed choices originally used for fitting. Here, choices were simulated 100 times for each participant using the mode of the derived per-participant parameter distribution. Simulated choice accuracy was averaged across simulations and evaluated against the observed choice data (Fig. 2c).

### Quantifying single-trial estimates

The modes of the per-participant posterior parameter distributions were selected to describe individual positive and negative learning rates (*α_Gain_*, *α_Loss_*) and relative reward sensitivity (*β*). In the learning phase, these per-participant parameter estimates were used to quantify Q-values and RPEs on each trial. Specifically, we quantified for each trial the value of the chosen option, the unchosen option and the difference between presented options (ΔValue). In the transfer phase, when participants did not receive feedback about their choices, we investigated how previously learned value related to pupil responses during value-based decisions. To do so, we selected the final Q-value estimates for each option (i.e. at the end of the learning phase) and used these values to quantify for each trial the value of the chosen and unchosen stimulus, given the individual sequences of choices, tte obtained single-trial variables were used as covariate regressors in a deconvolution analysis (described below), to investigate how they dynamically varied with trial-by-trial fluctuations in transient pupil responses in the learning and transfer phase.

### Pupillometry: preprocessing

The diameter of the pupil was recorded at a 1000Hz using anEyeLink 1000 Tower Mount (SR Research), tte eye-tracker was calibrated prior to each run. Blinks and saccades were detected using standard EyeLink software with default settings and Hedfpy, a Python package for preprocessing eye-tracking data. Periods of data loss during blinks were removed by linear interpolation, using an interpolation time window of 200ms before until 200ms after a blink. Blinks not identified by the manufacturer’s software were removed by linear interpolation around peaks in the rate of change of pupil size, using the same interpolation time window, tte interpolated pupil signal was band-pass filtered between 0.05Hz and 4Hz, using third-order Butterworth filters, z-scored per run, and resampled to 20Hz. As blinks and saccades have strong and relatively long-lasting effects on transient pupil size^61,62^, these influences were removed from the data, as follows. Blink and saccade regressors were created by convolving all blink and saccade events with their standard Impulse Response Function (IRF)^62–64^. ttese convolved regressors were used to estimate their responses in a General Linear Model (GLM), after which we used the residuals of this GLM for further analysis. For the subsequent deconvolution analysis, trials were removed in which participants made a saccade towards either of the two presented colored stimuli (i.e. saccades exceeding 3.3° visual angle away from fixation) to ensure that pupil responses were not affected by eye movements (percentage removed trials, mean=4.8%; SD=4.5%; range=0.0%-16.3%).

### Pupillometry: deconvolution analysis

#### Learning phase

Transient pupil responses were analyzed using FIRDeconvolution, a Python package used to perform Finite Impulse Response fits^65^. For the analysis of the learning phase, a design matrix was constructed that estimated pupil time courses of the following 3 transient event types: the onset of the choice options (start of the decision interval), choice (keypress) and feedback (auditory tone). Time courses of the onset of the options and feedback were estimated in the interval −0.5s. pre-event until 3.0s. post-event, tte time course of the choice was estimated in the interval −2.0s. pre-event until 1.5s post-event, as decision-related pupil dilation is predominantly driven prior to the behavioral report^5,35^. Further, the sustained drive of pupil size during the decision interval, defined as the time period from onset of the options until the choice, was estimated by a boxcar regressor, tte boxcar regressor expanded each trial’s RT (in samples) and was normalized by dividing the height of the boxcar by the mean RT of the regressor, ttis procedure ensured that the estimated IRF of all transient and sustained regressor types were comparable. Lastly, the design matrix included 2 stick regressors to estimate the pupil time course of the following nuisance events: the onset of the fixation dot and the offset of the options from the screen. Pupil time courses of both events were estimated −0.5s. pre-event until 3.0s. postevent. No intercept was added to the design matrix as ridge regression (described below) requires centered dependent and independent variables^66^. For the decision interval, we investigated how value beliefs about presented options affected pupil size by adding single-trial chosen and unchosen Q-value estimates as covariates to the design matrix, ttese Q-value estimates were also used as covariates for the feedback interval, to investigate how value expectations about a recent choice affected pupil size during feedback. Finally, we added single-trial RPE estimates to the design matrix to investigate how violations of choice beliefs affected the feedback-related response. All covariate regressors were z-scored per participant, to ensure unbiased across-subject comparisons of deconvolution beta weights.

#### Transfer phase

The design matrix for the deconvolution analysis of the transfer phase was identical to that of the learning phase, with two exceptions: (1) the pupil time course for feedback events was not estimated, as no feedback events occurred during this phase, (2) a stick regressor was included to investigate the effects of choice conflict on pupil responses. Choice conflict was determined the basis of experimental reinforcement value differences between the presented options, where trials were divided into three bins (10-20%; 30-40% and 50-60%), that corresponded to large, medium and small choice conflict between options.

### Pupillometry: ridge regression

We implemented the deconvolution analysis using cross-validated ridge regression, which allows one to find the general solution to a least-squares problem that would be unstable due to multicollinearity of regressors^66^. Ridge regression penalizes, or shrinks, regression coefficient weights towards zero to reduce the estimation variance on the coefficients:

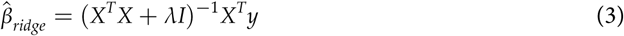

Here, *y* is the pupil time series signal and *X* is the design matrix consisting of a set of vectors that contain ones at all sample times relative to the event timings of which we estimated the pupil response, and zeros elsewhere, tte identity matrix, *I*, is multiplied by *λ* ≥ 0, a tuning parameter that controls the strength of the penalty term. If *λ* = 0, the linear regression solution is obtained, *λ* = ∞, *β̂_rifge_* 0. To obtain for each participant the optimal *λ* value, we applied cross validation on the pupil time series data. Here, the pupil data was divided into a training and test set. A weight matrix was obtained for each *λ* value (range = 0 ≤ *λ* ≤ 1), using the training set, and was used to predict the test set. ttis process was, repeated for 20 different selections of training and test sets, and the best *λ* value was selected based on, its prediction accuracy, tte resulting regression, *β̂_rifge_* contained the deconvolved pupil responses of all separate event types.

### Statistical comparisons

Nonparametric cluster-based permutation *t*-tests^67–69^ were used to test for significant regression coefficients and to correct for multiple comparisons over time. Briefly, for each time point of a time series signal, *t*-tests were performed on each set of across-subject coefficient values, tte cluster size was de! termined by the number of contiguous timepoints for which the *t*-test resulted in *P*<.05. tte observed; cluster size was then compared to a random permutation distribution of maximal cluster sizes: the i proportion of random clusters resulting in a larger size than the observed one determined the *P*-value, corrected for multiple comparisons.

To assess the effects of chosen and unchosen value covariates on pupil size across the decision interval, we summed each regressor’s coefficient values locked to the start (option onset) and locked to i the end of the decision interval (the moment of choice), while discarding their post-choice effects. We *i* normalised the summed regressor coefficient values by the number of samples they explained of the i pupil time series signal, tte resulting averaged, normalised regressor coefficient values were used in a repeated measures ANOVA to test for main and interaction effects on pupil size, both for the learning : and transfer phase.

Across-subject analyses of the relation between pupil responses and computational model parami eters were calculated using bootstraps^70^. We randomly drew with replacement 10,000 new pupil size -, model parameter estimate pairs which were used in the across-subject GLM. From the resulting boot, strapped regression coefficients, 68% confidence intervals were calculated using a percentile approach. *P*-values calculations were based on a two-sided hypothesis test, with the *P*-value being the fraction of the bootstrap distribution that fell below (or above) 0.

## Acknowledgements

We thank Lisa Roodermond and Lynn van den Berg for assistance in the data collection of the study, ttis work was supported by an ERC Advanced Grant ERC-2012-AdG-323413 to JT.

## Author contributions

JCS, SJ and TK designed the study. JCS collected and analysed the data. SJ and TK contributed novel methods. JCS and SJ wrote the first draft of the manuscript. JCS, SJ, TK and JT wrote the final manuscript.

